# A tool to shoot genes with massive air from a compressor (TSGMAC)

**DOI:** 10.64898/2026.03.24.713841

**Authors:** Daisuke Tsugama

**Affiliations:** Asian Research Center for Bioresource and Environmental Sciences (ARC-BRES), Graduate School of Agricultural and Life Sciences, The University of Tokyo, 1-1-1 Midori-cho, Nishi-tokyo-shi, Tokyo 188-0002, Japan

**Keywords:** fluorescent protein, onion, particle bombardment, rice, transformation

## Abstract

Particle bombardment systems are widely used for plant transformation, but commercial devices are expensive and rely on high-pressure helium gas. This study aimed to develop a cost-effective and helium gas-free alternative using an air duster gun connected to a commercial compressor. A nozzle (for DNA with transgenes), gold particles (as DNA carriers), nozzle-to-sample distance, and a method for coating gold particles with DNA were optimized to yield better transformation efficiency in targeting onion epidermal cells and rice calli. From the rice calli transformed with the newly developed system (a tool to shoot genes with massive air from a compressor: TSGMAC), stable transgenic plants could be obtained. TSGMAC offers a low-cost and helium gas-free solution for plant transformation and genome editing and can enhance accessibility to particle bombardment-based techniques.

## Introduction

Particle bombardment was originally developed by Sanford et al. (1987) and is widely used for introducing DNA, RNA, protein and other macromolecules into plant cells. Particle bombardment can be used for transforming species and tissues that are difficult to modify with DNA-transferring bacteria such as *Agrobacterium tumefaciens*, and for chloroplast and mitochondrial transformation (e.g., Boynton et al. 1988). Particle bombardment can also mediate direct delivery of genome-editing ribonucleoproteins (RNPs), which, for example, consist of Cas9 and its guide RNA, into plant cells, enabling DNA-free genome editing (Imai et al. 2020). Existing commercial particle bombardment systems such as PDS-1000/He (Bio-Rad, Hercules, CA, USA), Helios Gene Gun (Bio-Rad), and GDS-80 (Wealtec Bioscience Co., Ltd., Taipei, Taiwan) are well designed but expensive, requiring investments of tens of thousands of dollars. Moreover, all of these systems rely on high-pressure helium gas to accelerate particles, but helium is a limited resource and cannot be stably supplied. These factors hinder accessibility to those systems. As an alternative, a TSGAMAP (tool to shoot genes with a man-made air pressure) system was developed, offering a low-cost approach using compressed air from a manual pump for approximately $100 (Tsugama and Takano 2019). However, due to manual operation, its throughput and application are limited. Here, a new low-cost particle bombardment system with improved throughput that eliminates helium gas dependency is introduced.

## Materials and methods

### Preparation of the TSGMAC system

The TSGMAC system was assembled from the parts indicated in Fig. S1 and S2.

Gold particles with the average size of 0.6 μm were purchased from Bio-Rad (Hercules, CA, USA) and InBio Gold (Hurstbridge, Australia). Gold particles with the average size of either 1 μm were purchased from Bio-Rad. These particles were washed once with 99.5% (v/v) ethanol, once with 70% (v/v) ethanol, and three times with distilled water (DW). The gold particles were then resuspended in DW with the 60 mg/mL final concentration and stored at 4°C until used. To resuspend the gold particles in washing and splitting them, the particles were sonicated briefly (for ∼10 seconds) in an ultrasonic cleaning bath (As One Co., Osaka, Japan).

To coat the gold particles with DNA with spermidine and CaCl2, 2.5 M CaCl2 and 100 mM spermidine stock solutions were separately prepared and stored at 4°C. The spermidine stock was prepared within one week of its use for the TSGMAC.

To coat the gold particles with DNA with polyethylene glycol (PEG) and MgCl2, PEG with the average size of interest (Fig. S3) was dissolved in 160 mM MgCl2 in the concentration of interest. For example, for the “P2000/Mg” coating method in Fig. S3, a solution containing 50% (w/v) PEG2000 and 160 mM MgCl2 was prepared. These PEG and MgCl2 solutions were prepared within one week of their use for the TSGMAC.

The plasmids pBS-35SMCS-GFP (Tsugama et al. 2012), pBS-35SMCS-mCherry (Tsugama et al. 2013) and pCAMBIA1300 (Abcam, Cambridge, UK) were purified by the GenElute Plasmid Miniprep kit (Merck KGaA, Darmstadt, Germany) from the cells of the *Escherichia coli* strain DH5α that were transformed with them and that were cultured overnight at 37°C in the presence of 75 mg/L carbenicillin (for pBS-35SMCS-GFP and pBS-35SMCS-mCherry) or 50 mg/L kanamycin (for pCAMBIA1300). The DNA concentration of pBS-35SMCS-GFP and pBS-35SMCS-mCherry stock solutions was adjusted to 400 ng/μL, and that of the pCAMBIA1300 stock solution was adjusted to 150 ng/μL.

Bulbs of onion (*Allium cepa*) were purchased at a local supermarket within one month of their use for the TSGMAC and stored at 4°C. Seeds of rice (*Oryza sativa* ssp. *japonica* cv. Nipponbare) were purchased from Nouken Co. (Kyoto, Japan) and stored at 25°C. They were transformed as described below in the “Transient gene expression in onion scale leaf epidermis” and “Rice transformation” subsections.

### Transient gene expression in onion scale leaf epidermis

To coat the gold particles with either pBS-35SMCS-GFP or pBS-35SMCS-mCherry, a mixture in Table S1 was prepared using the components described above in the “Preparation of the TSGMAC system” subsection. The resulting mixture was sonicated for ∼10 seconds and incubated at 25°C for 5 minutes with regular resuspension of gold particles. The mixture was centrifuged at 10000× *g* for 5 seconds and the resulting supernatant was discarded. One hundred and fifty μL of 70% (v/v) ethanol was added, and the sample tube was inverted a few times to wash the gold particles. The tube was centrifuged at 10000 × *g* for 5 seconds and the resulting supernatants were discarded. One hundred and fifty μL of 99.5% (v/v) ethanol was added, and the sample tube was inverted a few times to wash the gold particles. The tube was centrifuged at 10000 × *g* for 5 seconds and the resulting supernatants were discarded. The gold particles were then resuspended in 99.5% (v/v) ethanol at the 50 mg/mL concentration and sonicated for 10 seconds immediately before they were bombarded.

Compressed air with 0.8 MPa was generated in the air compressor, and the pressure in the regulator was set as 0.3 MPa. Five μL of the gold particle solution was placed on the cylinder, the cylinder was inserted into the fitting with the de Laval nozzle, and these were fixed on the impact blow gun as shown in Fig. S1 and S2. A piece of the third outermost or an inner scale leaf of an onion bulb was manually held with the gun to make an intended distance (“nozzle-target distance”) between the adaxial side of the leaf and the nozzle edge, and then the gold particles were bombarded. A part of the epidermis including the site of the bombardment was isolated 12-20 hours after the bombardment was performed and subjected to the fluorescence microscopy to detect the signals of GFP and mCherry (Tsugama et al. 2012, 2013). GFP- and mCherry-positive cells were manually counted and used as the numbers of transformed cells shown in Fig. S4-S7. Some of the above settings are illustrated in Fig. S8.

### Rice transformation

Seeds of rice (*Oryza sativa* cv. Nipponbare) were purchased at Nouken Co. (Kyoto, Japan), surface-sterilized, sown on a callus-inducing medium (N6CIM, Table S2), and incubated at 28℃ in the darkness essentially as described previously (Toki et al., 1997). The resulting 20-40-day-old calli were placed on a holder of stainless steel sieve and autoclavable silicon rubber under an aseptic condition, fixed with tweezers and stainless steel mesh with the size 20 mesh/inch, and shot with the modified “P3350/Mg” condition (Table S1, bottom, and Fig. S3) and the 0.2 MPa pressure in the regulator. Some of these settings are illustrated in Fig. S9. The bombarded calli were incubated on N6CIM at 28℃ in the darkness for three days and transferred to N6CIM supplemented with 30 mg/L hygromycin B (Fujifilm Wako, Osaka, Japan) (N6CIMH, Table S2). The calli were then incubated at 28℃ in the darkness for 14 days there, transferred to new N6CIMH, and incubated in the darkness for 14 days. The calli growing there were transferred to shoot induction media (SIM, Table S2) and incubated at 28℃ under the 12-h light/12-h darkness photoperiod until shoots emerged. The shoot tissues were transferred to root induction media (RIM, Table S2) when they were 0.5-1 cm in length. The tissues were incubated at 28℃ under the 12-h light/12-h darkness photoperiod until roots emerged and the shoots became ∼5 cm in length. The resulting seedlings were transferred to a soil mixture consisting of akadama, vermiculite, charcoal, slow-release, and quick-release fertilizers (“Shiba no Metsuchi Tokotsuchi”, Tachikawa Heiwa Nouen, Kanuma, Japan) in plastic pots with holes, and further grown with regular watering essentially as previously described (Banakar and Wang 2020). GFP signals were detected with an epifluorescence microscope (BX51, Olympus, Hachioji, Japan) equipped with a CCD camera (DP73, Olympus) and a fluorescence mirror unit (U-MWIB2, Olympus) occasionally when the calli were incubated on the N6CIMH and when the plants were in the 5- or 6-leaf stage on the soil. Genomic DNA was prepared with the DNeasy Plant Mini kit (Qiagen, Hilden, Germany) from leaf tips of approximately 3 cm in length of the individual plants growing on the soil when they were in the 5- or 6-leaf stage. The resulting DNA was used as templates for the PCR to confirm the presence of the transgenes in the putative transgenic plants. The primers used are listed in Table S3.

## Results and discussion

The new particle bombardment system, called TSGMAC (tool to shoot genes with massive air from a compressor), consists of an impact blow (air duster) gun, a de Laval nozzle, a pressure regulator and an air compressor, which cost approximately $300 at minimum. Gold particles coated with DNA or other molecules are placed on a cut pipette tip (“cylinder”), which is placed in a nozzle fitting (Fig. 1 and S1). The particles are released toward a target tissue by a high-pressure air from the compressor and accelerated by the aerodynamic effect of the de Laval nozzle (Fig. S2). The components and methods for the operation of this system were optimized for onion scale epidermis and rice calli as target tissues (Table S1 and S2; Fig. S3-S9). This resulted in yielding dozens of cells per bombardment in both the onion epidermis (Fig. 2a) and rice calli (Fig. 2b, left panel). Co-bombardment of pCAMBIA1300 (a vector with the hygromycin resistance gene *HPT* downstream of the 35S promoter) and pBS-35SMCS-GFP (a vector with *GFP* downstream of the 35S promoter), into rice calli yielded hygromycin-resistant and GFP-positive calli (Fig. 2b, right panel). From these calli, transgenic plants with *HPT* and *GFP* could be regenerated (Fig. 2c and d). These results indicate that TSGMAC can be used for both transient gene expression in multiple cells with multiple constructs and obtaining stable transgenic plants. TSGMAC offers a low-cost and helium gas-free solution for plant transformation and genome editing and can enhance accessibility to particle bombardment-based techniques.

**Figure 1.**
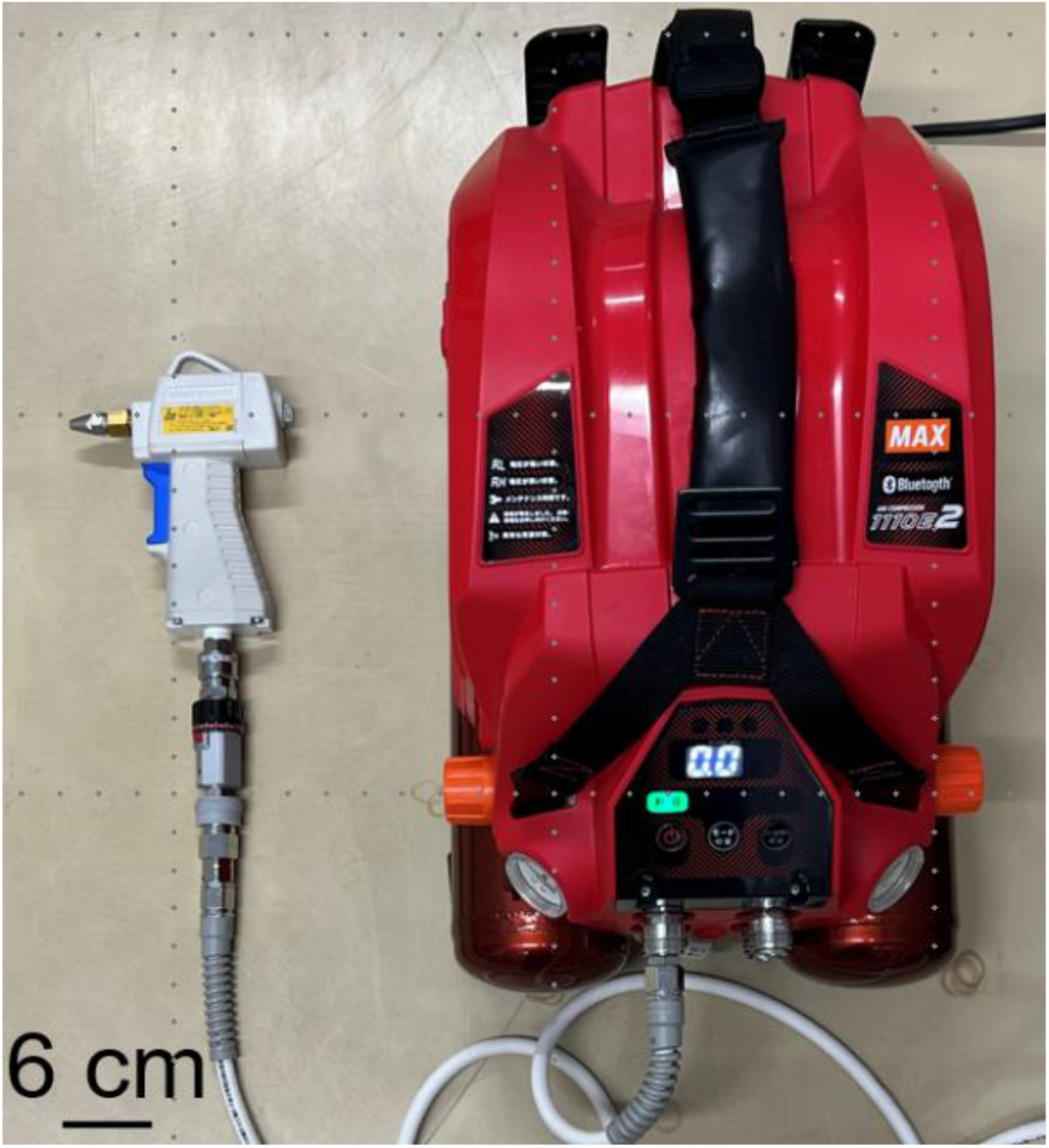
Appearance of TSGMAC. Details about the components of TSGMAC are given in Fig. S1.

**Figure 2.**
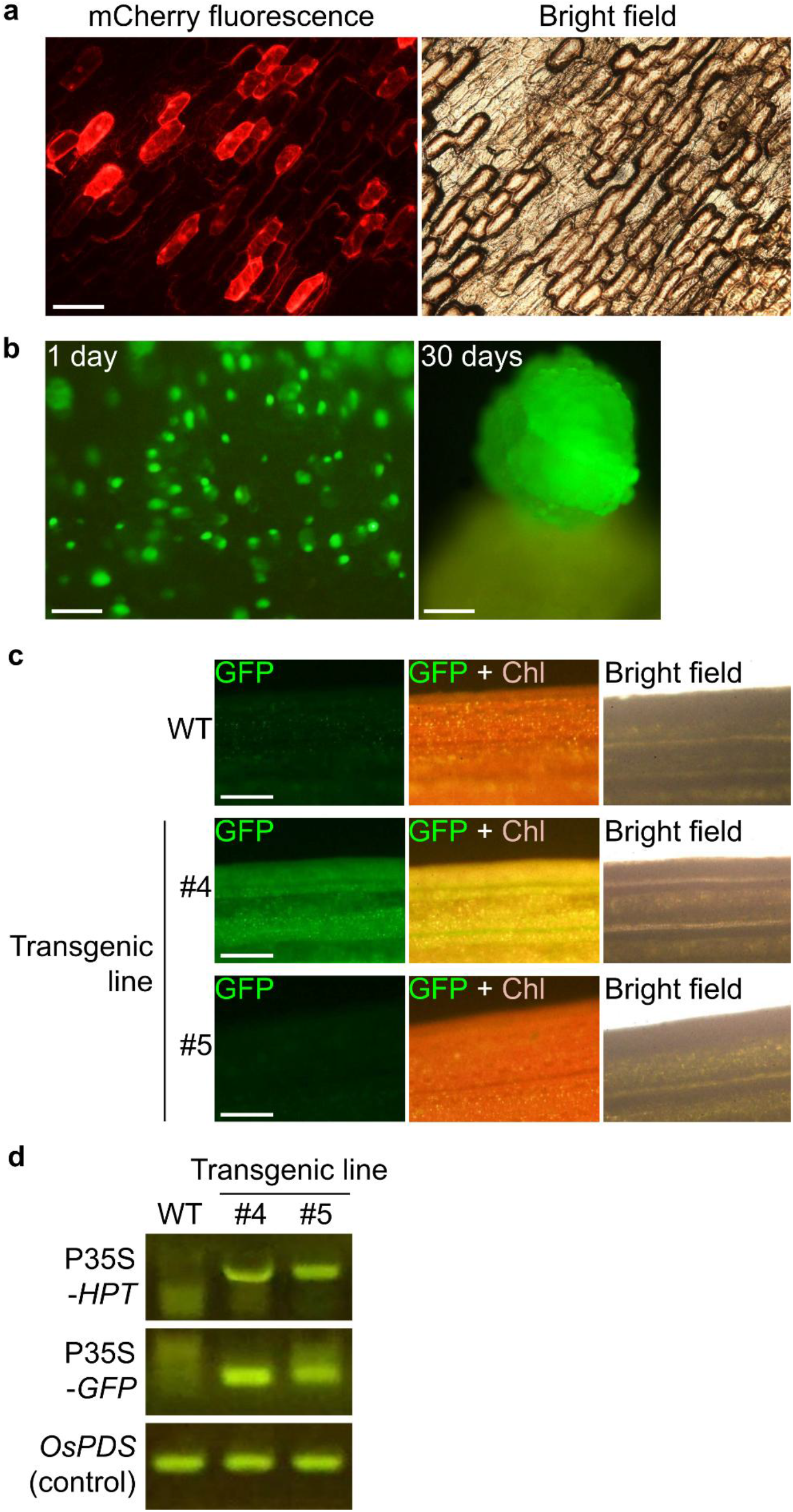
Use of TSGMAC for transient gene expression in plant cells and for generating stable transgenic plants. (a) Transient expression of mCherry in onion epidermal cells. The mCherry expression construct pBS-35SMCS-mCherry (Tsugama et al. 2013) was introduced into the onion cells by the TSGMAC as described in Supplementary Methods under the condition highlighted in Supplementary Fig. S3. Scale bar = 300 μm. (b) Expression of GFP in cells of rice calli. The GFP expression construct pBS-35SMCS-GFP (Tsugama et al. 2012) and the *HPT*-containing vector pCAMBIA1300 (Abcam, Cambridge, UK) were introduced into the cells of rice calli by the TSGMAC as described in Supplementary Methods under the condition highlighted in Supplementary Fig. S3. GFP signals detected in the calli one day and 30 days after the bombardment are presented in the left and right panels, respectively. Scale bars = 150 μm in the left panel and 300 μm in the right. (c) GFP signals in leaves of the wild type and stable rice transgenic lines #4 and #5. Young leaves of the plants at the 5- or 6-leaf stage were used for the fluorescence microscopy. GFP signals in the line #5 were undetectable. Chl: chlorophyll autofluorescence. Scale bars = 300 μm. (d) Genomic PCR analysis of the transgenes. P35S-*HPT*: 35S promoter-*HPT* region from pCAMBIA1300; P35S-*GFP*: 35S-*GFP* region from pBS-35SMCS-GFP.

## Acknowledgments

This study was supported by JSPS (Japan Society for the Promotion of Science) KAKENHI Grant [grant numbers: 23K19293 and 25K09059].

## Statements and Declarations

### Author Contributions

DT designed and performed the experiments and data analysis and wrote the manuscript.

### Data Availability

All data generated or analyzed during this study are included in this published article and its supplementary information file.

