## Supplementary materials for "A tool to shoot genes with massive air from a compressor (TSGMAC)"

Daisuke Tsugama

Asian Research Center for Bioresource and Environmental Sciences (ARC-BRES), Graduate School of Agricultural and Life Sciences, The University of Tokyo
1-1-1 Midori-cho, Nishi-tokyo-shi, Tokyo 188-0002, Japan


**Table S1. Mixtures for coating gold particles with DNA for the TSGMAC (for four bombardments)**

- **Spermidine- and $CaCl_2$-based mixture for onion scale leaves**

| Item | Amount (μL) | Final concentration |
|---|---|---|
| 60 mg/mL gold particle suspension | 16 | 24 μg/μL (mg/mL) |
| DW | 0.6 | - |
| 400 ng/μL DNA | 1 | 10 ng/μL |
| 2.5 M $CaCl_2$ | 16 | 1 M |
| 100 mM spermidine | 6.4 | 16 mM |

- **PEG- and $MgCl_2$-based mixture for onion scale leaves**

| Item | Amount (μL) | Final concentration |
|---|---|---|
| 60 mg/mL gold particle suspension | 16 | 24 μg/μL (mg/mL) |
| DW | 3 | - |
| 400 ng/μL DNA | 1 | 10 ng/μL |
| PEG with $MgCl_2$ or $CaCl_2$ | 20 | See Fig. S3 |

- **PEG- and $MgCl_2$-based mixture for rice**

| Item | Amount (μL) | Final concentration |
|---|---|---|
| 60 mg/mL gold particle suspension | 16 | 24 μg/μL (mg/mL) |
| 150 ng/μL pCAMBIA1300 | 7 | 21 ng/μL |
| 400 ng/μL DNA | 2 | 16 ng/μL |
| 50% (w/v) PEG 3350 in 160 mM $MgCl_2$ | 25 | 25% (w/v) PEG 3350; 80 mM $MgCl_2$ ("P3350/Mg" in Fig. S3) |

### Table S2. Media used for rice transformation

| Medium | Component | Final concentration | Manufacturer | Note |
|---|---|---|---|---|
| N6CIM | Chu N6 salt mixture | 4.1 g/L (1x) | Fujifilm Wako, 391-02021 | Stock: 4.1 g/L in distilled water (DW), autoclaved |
| N6CIM | Sucrose | 30 g/L | Fujifilm Wako, 190-00013 | |
| N6CIM | Acid hydrolysate of casein (Casamino acid) | 0.3 g/L | Solabia Biokar Diagnostics, A1404HA | |
| N6CIM | L-proline | 2.8 g/L | Fujifilm Wako, 161-04602 | |
| N6CIM | 2,4-dichlorophenoxyacetic acid (2,4-D) | 2 mg/L | Fujifilm Wako, 040-18532 | Stock: 20 mg/mL in ethanol; Added before autoclaving |
| N6CIM | Plant Preservative Mixture (PPM) | 1 mL/L (0.1 % (v/v)) | Plant Cell Technology, 100 PPM (Nacalai Tesque, 26062-84) | Added before autoclaving |
| N6CIM | Potassium hydroxide (KOH) | Approx.0.95 mL of 1 M stock/L (to make pH 5.8) | Fujifilm Wako, 168-21815 | Stock: 1 M in DW |
| N6CIM | Gellan Gum | 4 g/L | Fujifilm Wako, 075-03075 | |
| N6CIM | N6 vitamin solution (1000x) | 1 mL/L (0.1% (v/v), 1x) | Glycine: Fujifilm Wako, 077-00735<br>Thiamin hydrochloride: Fujifilm Wako, 201-00852<br>Pyridoxine hydrochloride: Fujifilm Wako, 163-05402<br>Nicotinic acid: Fujifilm Wako, 142-01232 | Stock: 2 g/L glycine, 1 g/L thiamin HCl, 0.5 g/L pyridoxine HCl and 0.5 g/L nicotinic acid in DW, autoclaved<br>The stock is added to the medium after autoclaving |
| N6CIMH | Hygromycin B | 30 mg/L | Fujifilm Wako, 089-06151 | Stock: 30 mg/mL in 70% (v/v) ethanol |
| SIM | Murashige and Skoog basal salt mixture | 4.6 g/L (1x) | Fujifilm Wako, 392-00591 | Stock: 4.6 g/L in DW, autoclaved |
| SIM | Maltose monohydrate | 40 g/L | Fujifilm Wako, 130-00615 | |
| SIM | Acid hydrolysate of casein (Casamino acid) | 1 g/L | Solabia Biokar Diagnostics, A1404HA | |
| SIM | Plant Preservative Mixture (PPM) | 1 mL/L (0.1% (v/v)) | Plant Cell Technology, 100 PPM (Nacalai Tesque, 26062-84) | Added before autoclaving |
| SIM | 6-benzyladenine (BA) | 3 mg/L | Fujifilm Wako, 026-07623 | Stock: 3 mg/mL in DMSO; Added before autoclaving |
| SIM | 1-naphthylacetic acid | 0.1 mg/L | Tokyo Chemical Industry, N0005 | Stock: 1 mg/mL in DMSO; Added bofore autoclaving |
| SIM | Potassium hydroxide (KOH) | Approx.0.8 mL of 1 M stock/L (to make pH 5.8) | Fujifilm Wako, 168-21815 | Stock: 1 M in DW |
| SIM | Agarose S | 10 g/L | Nippon Gene (Fujifilm Wako), 312-01193 | |
| SIM | Gamborg's vitamin solution (1000x) | 1 mL/L (0.1% (v/v), 1x) | Merck, G1019-50ML | Added after autoclaving |
| SIM | Antibiotic such as hygromycin and G-418 | 0 | | No antibiotic is used |
| RIM | Murashige and Skoog basal salt mixture | 2.3 g/L (0.5x) | Fujifilm Wako, 392-00591 | Stock: 4.6 g/L in DW, autoclaved |
| RIM | Maltose monohydrate | 10 g/L | Fujifilm Wako, 130-00615 | |
| RIM | Gamborg's vitamin solution (1000x) | 1 mL/L (0.1% (v/v), 1x) | Merck, G1019-50ML | Added after autoclaving |
| RIM | Agarose S | 8 g/L | Nippon Gene (Fujifilm Wako), 312-01193 | |

#### Table S3. Primers used for the genomic PCR for detecting transgenes in rice

| name | sequence (5' -> 3') | note |
|---|---|---|
| L35S-600-24_F | GGCCATCGTTGAAGATGCCTCTGC | Anneals to a long version of the 35S promoter in pBS-35SMCS-GFP |
| GFPin80_R | GGACACGCTGAACTTGTGGCCG | Used with L35S-600-24_F |
| 2x35S-600-24_F | GGCTATCGTTCAAGATGCCTCTGC | Anneals to the double 35S promoter in pCAMBIA1300 |
| HPTin500_R | TGCGCGACGGACGCACTGACGGTGTCGTCCATC | Used with 2x35S-600-24_F |
| gOsPDS_610_F | TTTGGGTGGAAAGGTTTTACTCTTATG | For a control |
| gOsPDS_1330_R | CCAAACAAGTTCTGTATGTTGGGATAA | For a control |

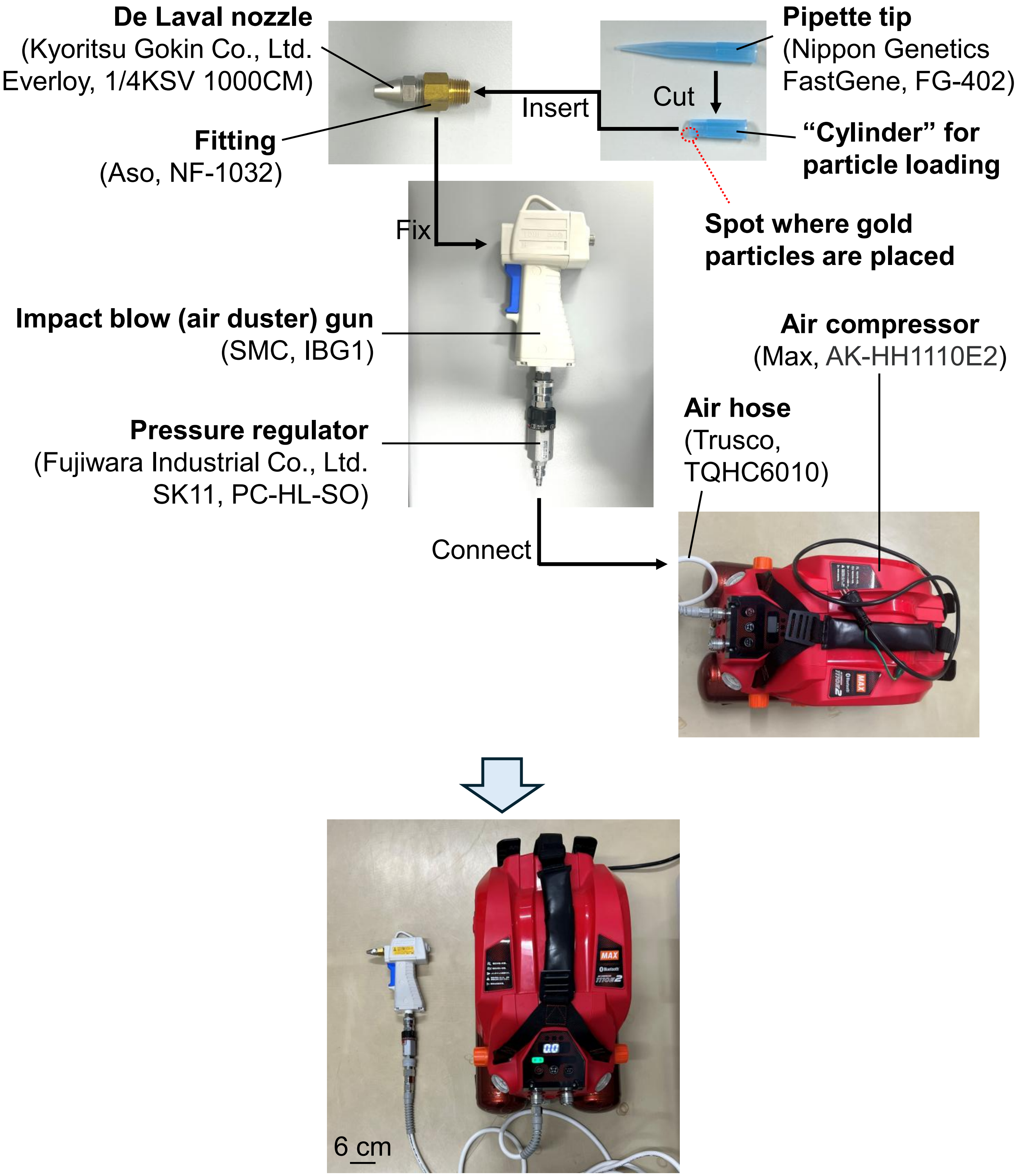

**Figure S1. The TSGMAC system.** The specific products used to generate the system are shown in the parentheses. All of these products except the impact blow gun IBG1 are compatible with 1 MPa or higher gas pressure, but such high pressure is not necessary at least for onion epidermis or rice calli. The bottom image is the same as Fig. 1.

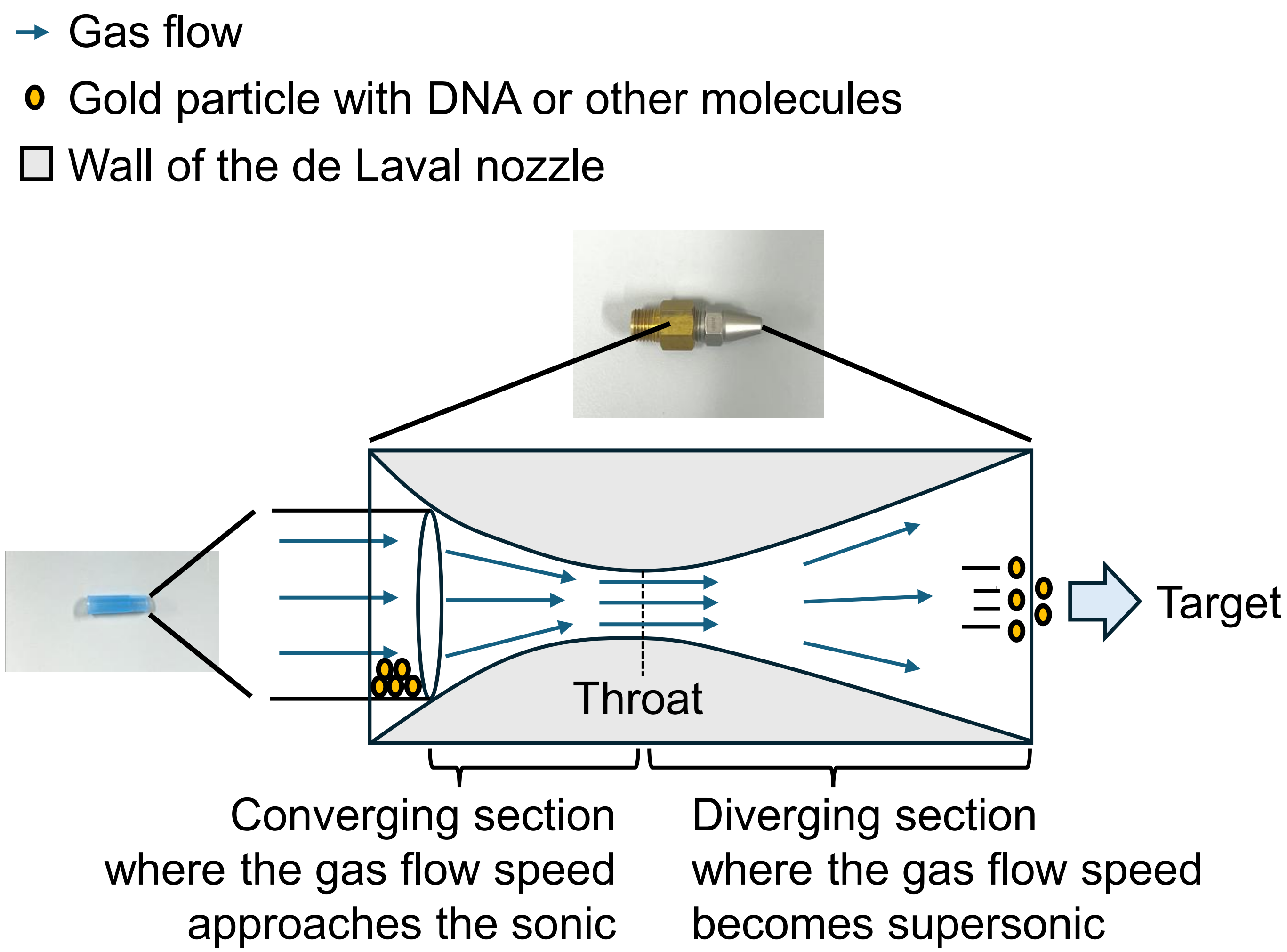

**Figure S2. Gas and particle dynamics in the TSGMAC.**

**Gold particle size**
- **0.6 um**
- 1.0 um

**Nozzle-target distance**
- 0.5 cm
- **1.5 cm**

**Cylinder aperture**

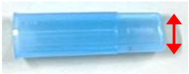

- 2 mm
- 2.5 mm
- 3 mm
- **3.5 mm**

**PEG 400 concentration (% (v/v))**
- 12
- 24
- **42**

**PEG 2000 concentration (% (w/v))**
- 6
- 12
- **25**

**PEG 3350 concentration (% (w/v))**
- 6
- 12
- **25**

**PEG 6000 concentration (% (w/v))**
- 6
- 12
- **25**

**PEG 20000 concentration (% (w/v))**
- 3
- **6**
- 12.5

**PEG 500000 concentration (% (w/v))**
- 0.02
- 0.1
- **0.25**
- 0.5
- 1

**$MgCl_2$ and $CaCl_2$ concentration** (with 10 ng/μL DNA and 24 μg/μL gold particle)
- 25% (w/v) PEG 3350 alone
- 25% (w/v) PEG 3350 + 20 mM $MgCl_2$ ("P3350/Mg")
- **25% (w/v) PEG 3350 + 80 mM $MgCl_2$ ("P3350/Mg")**
- 25% (w/v) PEG 3350 + 250 mM $MgCl_2$ ("P3350/Mg")

**Coating method** (with 10 ng/μL DNA and 24 μg/μL gold particle)
- 16 mM spermidine + 1 M $CaCl_2$ ("Spd/Ca")
- 42% (v/v) PEG 400 + 80 mM $MgCl_2$ ("P400/Mg")
- 25% (w/v) PEG 2000 + 80 mM $MgCl_2$ ("P2000/Mg")
- **25% (w/v) PEG 3350 + 80 mM $MgCl_2$ ("P3350/Mg")**
- 25% (w/v) PEG 6000 + 80 mM $MgCl_2$ ("P6000/Mg")
- 6% (w/v) PEG 20000 + 80 mM $MgCl_2$ ("P20000/Mg")
- 0.25% (w/v) PEG 500000 + 80 mM $MgCl_2$ ("P500000/Mg")

**Figure S3. An overview of the components and methods optimized for the operation of the TSGMAC.** The components and the methods that gave the largest numbers of transformed cells in the optimization experiments using onion scale leaves were used to compare the methods for coating gold particles with DNA and are indicated as bold and underlined texts. All experiments with these components and methods were performed with 10 ng/μL DNA (either pBS-35SMCS-GFP or pBS-35SMCS-mCherry) and 24 μg/μL gold particles. All the experiments with PEG (polyethylene glycol) were performed with 80 mM $MgCl_2$ as indicated in the bottom panel (for "Coating method"). The method names such as "Spd/Ca" in the bottom panel correspond to the names shown in Fig. S6. The highlighted condition (i.e., "P3350/Mg") was used thereafter for routine experiments.

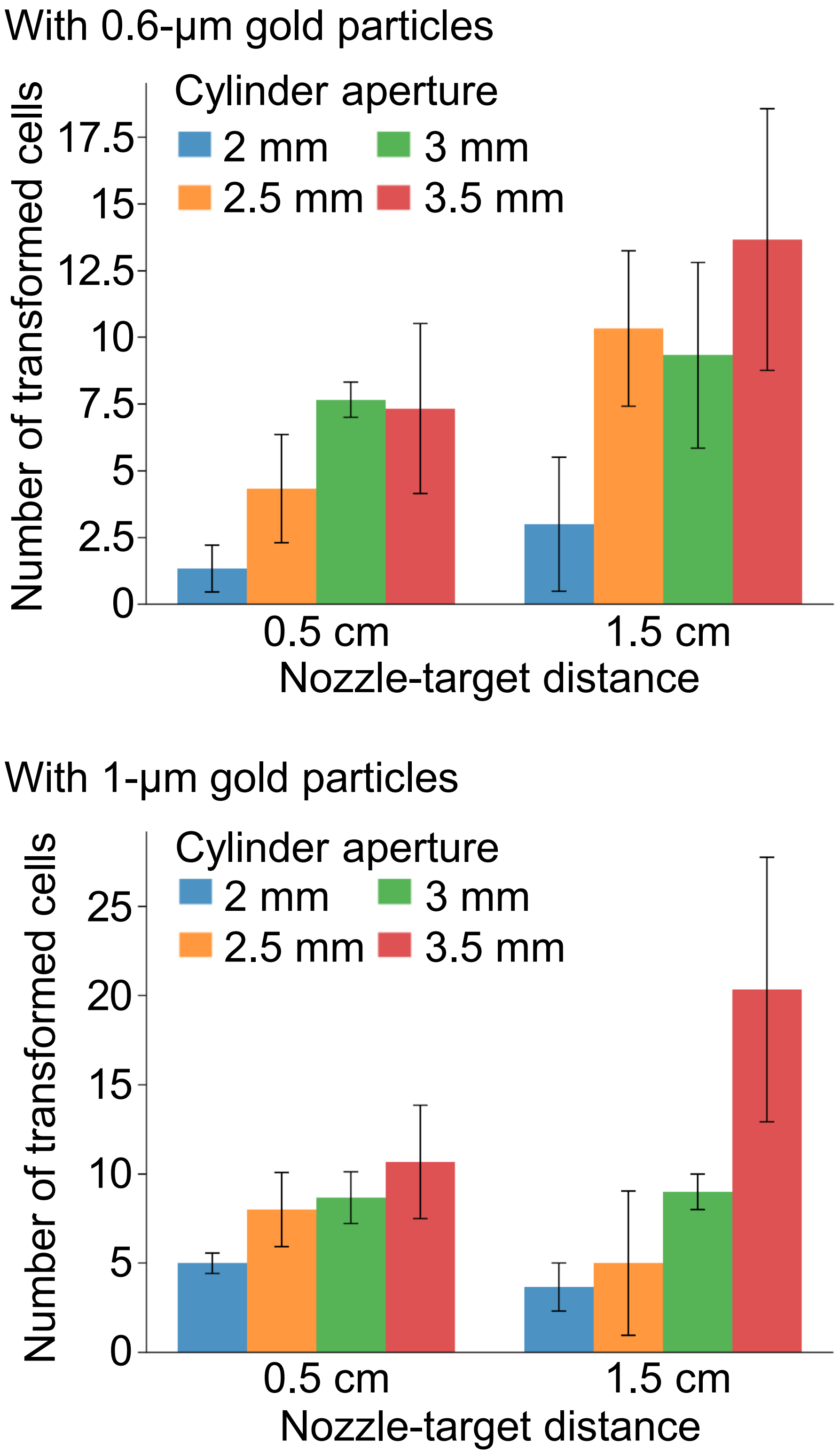

**Figure S4. Effects of the gold particle size, the cylinder aperture and the nozzle-target distance in operating the TSGMAC on the number of the transformed onion epidermal cells.** Either 0.6-μm gold particles (Bio-Rad) or 1-μm gold particles (Bio-Rad) were coated with pBS-35SMCS-GFP and bombarded at onion scale leaf epidermis by the TSGMAC. The numbers of the GFP-positive cells were counted under a fluorescence microscope 12-20 hours after they were transformed by the TSGMAC. Experiments were performed four times for each condition and means ± SD from them are presented.

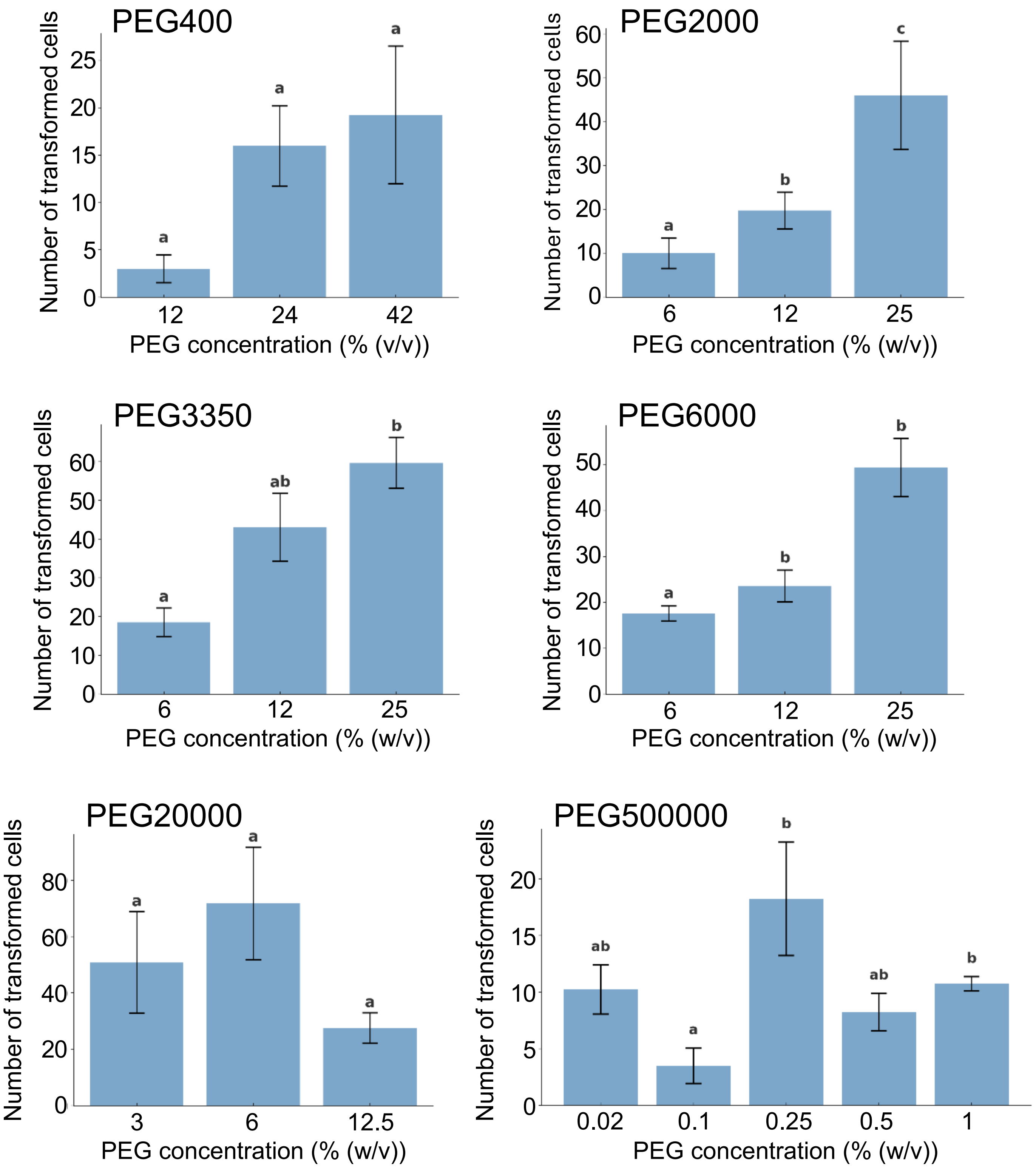

**Figure S5. Effects of the PEG size and concentration in operating the TSGMAC on the number of the transformed onion epidermal cells.** The 0.6-µm gold particles (InBio Gold) were coated with pBS-35SMCS-GFP in 80 mM $MgCl_2$ and the indicated concentration of PEG and bombarded at onion scale leaf epidermis by the TSGMAC. The numbers of the GFP-positive cells were counted under a fluorescence microscope 12-20 hours after they were transformed by the TSGMAC. Experiments were performed four times for each condition and means ± SD from them are presented. Data with different letters are significantly different ($P < 0.05$) according to the Tukey-Kramer test.

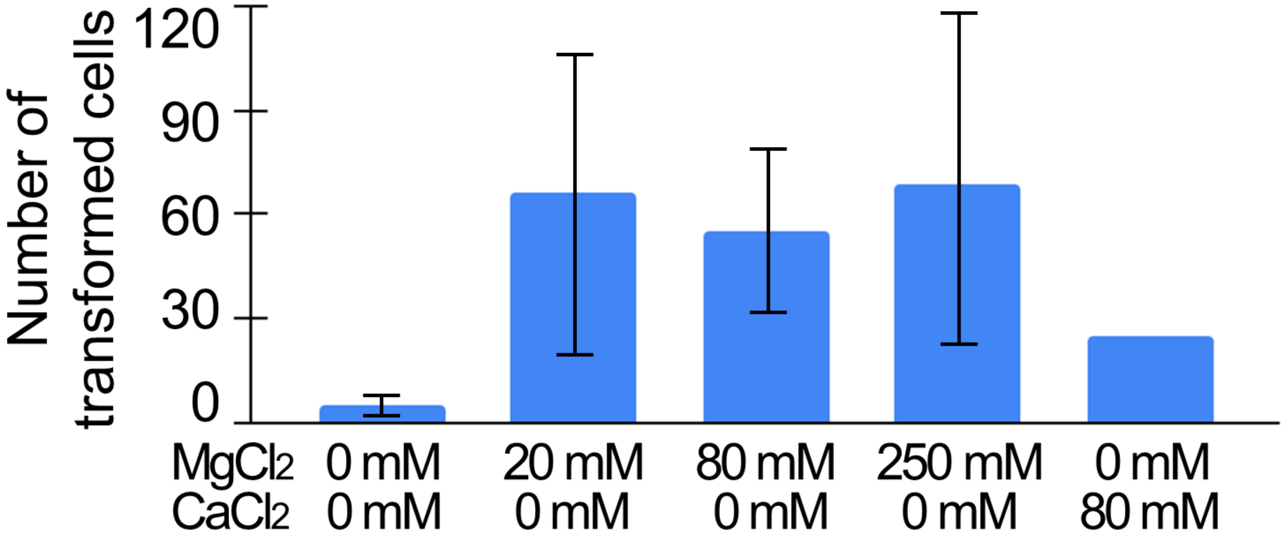

**Figure S6. Effects of the $MgCl_2$ and $CaCl_2$ concentration in operating the TSGMAC on the number of the transformed onion epidermal cells.** The 0.6-µm gold particles (InBio Gold) were coated with pBS-35SMCS-GFP in the solution containing 25% (w/v) PEG3350 and either $MgCl_2$ and $CaCl_2$ with the indicated concentration, and bombarded at onion scale leaf epidermis by the TSGMAC. The numbers of the GFP-positive cells were counted under a fluorescence microscope 12-20 hours after they were transformed by the TSGMAC. Experiments were performed five times for each condition and means ± SD from them are presented. P-value was smaller than 0.003 in the Kruskal-Wallis test.

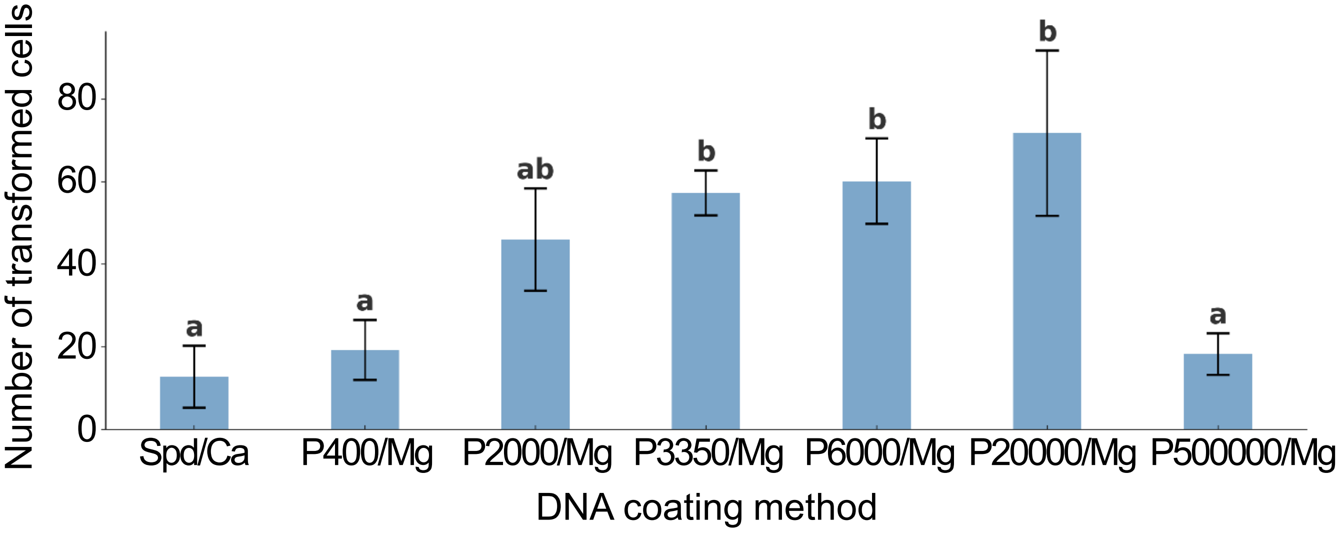

**Figure S7. A comparison of the methods for coating gold particles with DNA for the TSGMAC.** The 0.6-μm gold particles (InBio Gold) were coated with pBS-35SMCS-mCherry by the indicated methods. The numbers of the mCherry-positive cells were counted under a fluorescence microscope 12-20 hours after they were transformed by the TSGMAC. Experiments were performed four times for each condition and means ± SD from them are presented. Data with different letters are significantly different ($P$ < 0.05) according to the Tukey-Kramer test.

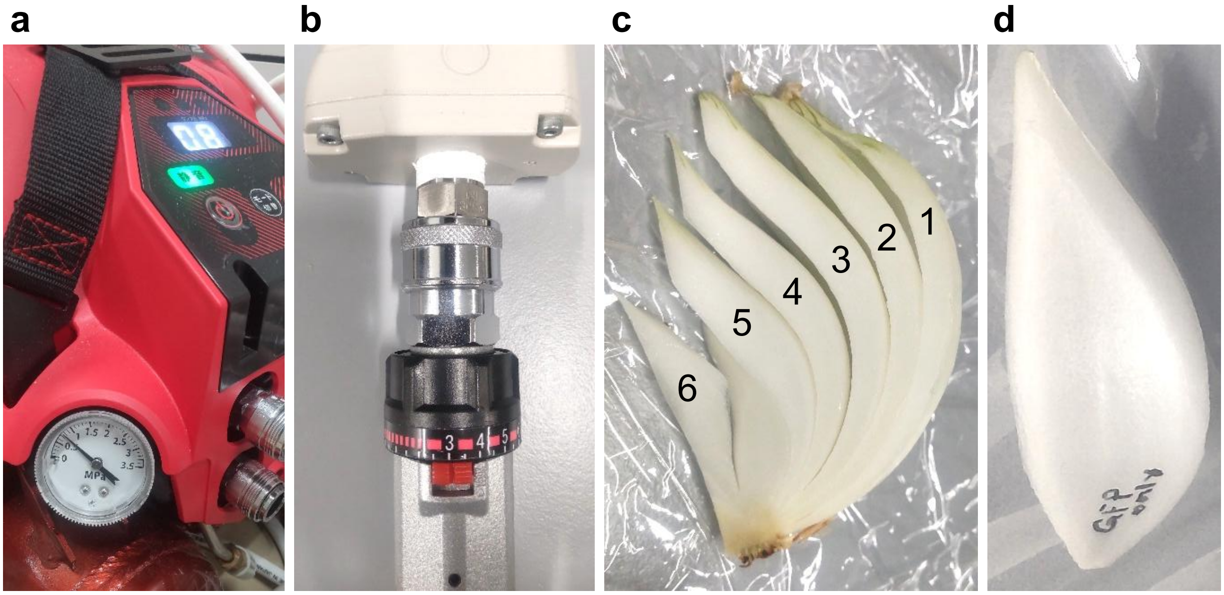

**Figure S8. TSGMAC settings for onion scale leaf epidermis.** (a) The pressure in the air compressor. (b) The pressure in the regulator. (c) The scale leaves used as the targets for the TSGMAC. The scale leaf pieces 3-5 in the figure could be used as such. (d) The post-bombardment appearance of a scale leaf. Only a few traces of gold particles (and the air-derived damages) are visible even when many transformed cells are obtained.

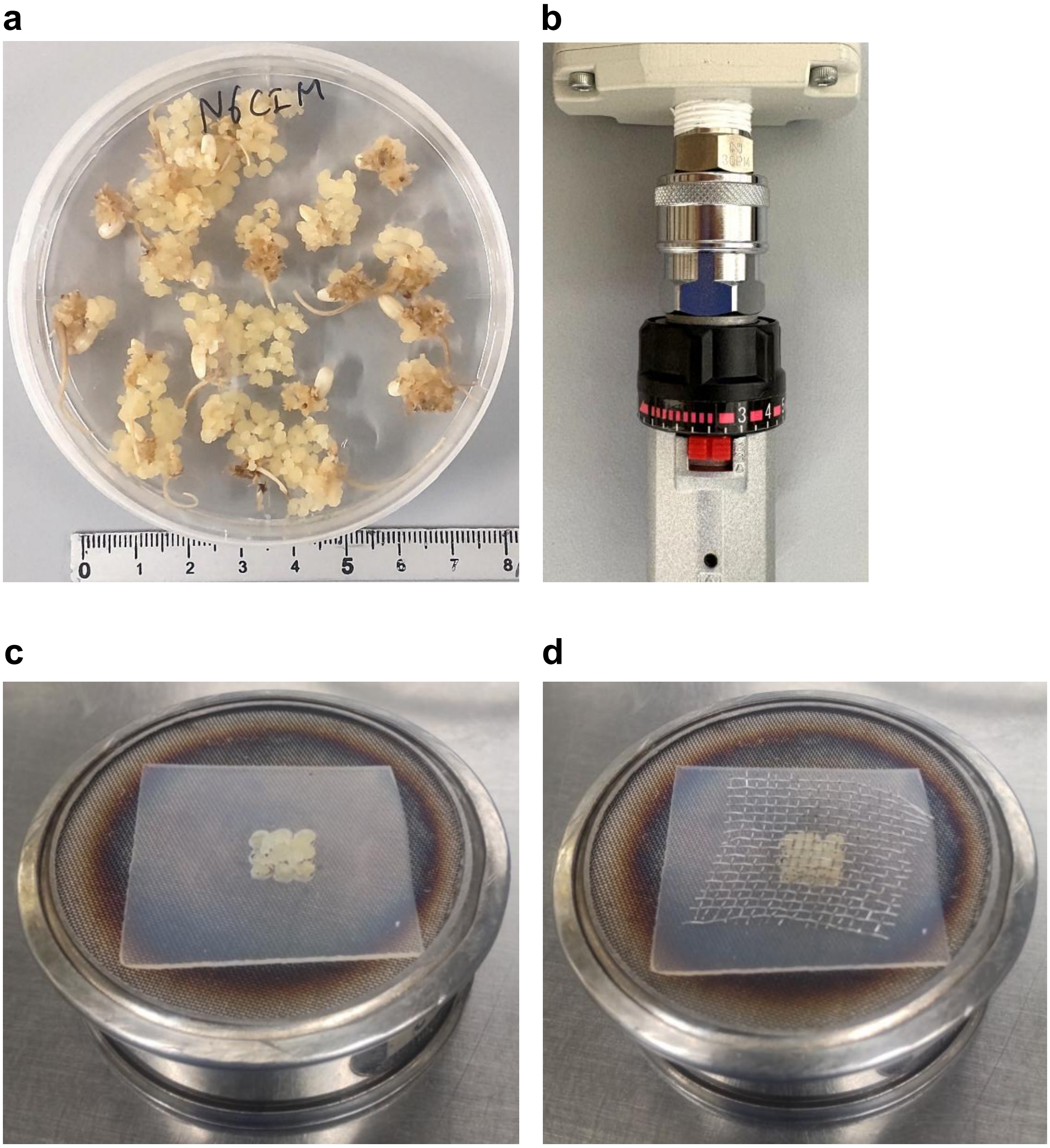

**Figure S9. TSGMAC settings for rice calli.** (a) Appearance of 35-day-old calli grown on the N6-based callus induction medium (N6CIM) and used as the TSGMAC-mediated transformation. (b) The pressure in the regulator. (c) The calli placed on a holder of stainless steel sieve and autoclavable silicon rubber under an aseptic condition. (d) The calli between the holder and stainless steel mesh. The mesh was pressed with tweezers to fix the calli immediately before they were bombarded.
